## Supplementary file for "Quantitative MRI reveals differences in striatal myelin in children with DLD"

***Supplementary file 1a.*** TD > DLD differences in MTsat maps. Nonparametric randomisation analysis with threshold-free cluster enhancement was used to compare groups. A whole brain corrected threshold of p<.05 was used.

| **Brain Area** | **X** | **Y** | **Z** | **1-p** | **Voxels** |
| --- | --- | --- | --- | --- | --- |
| ***MTsat*** | | | | | |
| ***Left temporal pole*** | -50 | 14 | -39 | 0.957 | 319 |
| ***Left Inferior Frontal Gyrus, pars opercularis*** | -50 | 13 | 6 | 0.962 | 194 |
| ***Left insula and opercular cortex*** |  |  |  | 0.975 | 5817 |
| *Long gyrus of the insular cortex* | -41 | -8 | 2 |  |  |
| *Central opercular cortex* | -50 | -8 | 10 |  |  |
| *Subcentral gyrus* | -60 | -8 | 8 |  |  |
| *Subcentral gyrus* | -60 | -12 | 20 |  |  |
| *Superior temporal cortex* | -56 | -13 | 0 |  |  |
| *Transverse temporal gyrus (Heschl’s)* | -54 | -19 | 9 |  |  |
| ***Left and Right Caudate nucleus*** |  |  |  | 0.992 | 11239 |
| *Left caudate nucleus (body)* | -14 | -10 | 24 |  |  |
| ***Left planum temporale*** | -56 | -37 | 17 | 0.967 | 414 |
| ***Right posterior parietal cortex*** | 44 | -69 | 49 | 0.967 | 768 |
| ***Left dorsal occipital and parietal cortex*** |  |  |  | 0.993 | 25842 |
| *Posterior middle temporal cortex* | -62 | -55 | -3 |  |  |
| *Intra-parietal sulcus (posterior part)* | -31 | -74 | 50 |  |  |
| *Parieto-occipital sulcus* | -8 | -82 | 42 |  |  |
| ***Right dorsal occipital and parietal cortex*** | 38 | -87 | 29 | 0.976 | 5414 |
| ***Left occipital pole*** | -19 | -94 | 17 | 0.964 | 600 |

***Supplementary file 1b****.* TD > DLD conjoint differences in R1 and MTsat

| **Brain Area** | **X** | **Y** | **Z** | **Voxels** |
| --- | --- | --- | --- | --- |
| *Right anterior cingulate cortex* | 1 | 14 | 39 | 307 |
| *Left inferior frontal gyrus, pars opercularis* | -53 | 11 | 0 | 195 |
| *Right thalamus* | 2 | -6 | -12 | 10637 |
| *Left insular cortex* | -34 | -13 | -7 | 1 |
| *Left planum polare* | -42 | -15 | -5 | 4632 |
| *Left planum temporale* | -53 | -38 | 13 | 419 |
| *Left middle temporal gyrus, temporooccipital part* | -64 | -56 | -8 | 1281 |
| *Right lateral occipital cortex, superior division* | 16 | -57 | 55 | 422 |
| *Left lateral occipital cortex, inferior division* | -49 | -66 | 7 | 1020 |
| *Left lateral occipital cortex, inferior division* | -51 | -70 | -4 | 225 |
| *Left lateral occipital cortex, superior division* | -54 | -71 | 21 | 120 |

***Supplementary file 1c.*** Differences in age and language scores between the selected and excluded children who were typically-developing (TD) or had developmental language disorder (DLD)

|  | **TD Selected** | **TD Unselected** | **p-value** | **DLD Selected** | **DLD Unselected** | **p-value** |
| --- | --- | --- | --- | --- | --- | --- |
| Age | 12.41 (1.62) | 11.69 (1.58) | 0.12 | 12.48  (1.80) | 11.39 (1.54) | **0.03** |
| ***Language tests*** | | | | | | |
| TROG-E | 105.23 (8.4) | 107.13 (7.04) | 0.38 | 81.33 (11.25) | 83.65 (14.37) | 0.57 |
| CELF Recalling Sentences | 11.85 (2.16) | 12.0 (2.65) | 0.84 | 5.03 (2.60) | 4.56 (2.73) | 0.55 |
| ROWVPT | 128.62 (16.48) | 133.31 (11.6) | 0.21 | 101.61 (15.83) | 98.72 (15.81) | 0.54 |
| EOWPVT | 117.36 (14.64) | 124.19 (15.32) | 0.13 | 91.61 (10.45) | 92.06 (15.43) | 0.91 |
| ERRNI Comprehension | 105.89 (13.77) | 111.62 (10.8) | 0.09 | 93.52 (14.63) | 94.17 (16.02) | 0.89 |
| ERRNI Initial Recall | 100.7 (11.86) | 100.38 (12.29) | 0.93 | 82.67 (12.62) | 85.72 (13.48) | 0.43 |
| ERRNI Delayed Recall | 105.55 (11.64) | 103.88 (8.43) | 0.53 | 85.76 (11.81) | 86.06 (14.2) | 0.94 |
